# An expectation-maximization-like algorithm enables accurate ecological modeling using longitudinal metagenome sequencing data

**DOI:** 10.1101/288803

**Authors:** Chenhao Li, Kern Rei Chng, Tamar V. Av-Shalom, Lisa Tucker-Kellogg, Niranjan Nagarajan

## Abstract

The dynamics of microbial communities is driven by a range of interactions from symbiosis to predator-prey relationships, the majority of which are poorly understood. With the increasing availability of high-throughput metagenome profiling data, it is now conceivable to directly learn ecological models that explicitly define microbial interactions and explain community dynamics. The applicability of these approaches is severely limited by the lack of accurate biomass and absolute density measurements. We present a new computational approach that resolves this key limitation in the inference of generalised Lotka-Volterra models (gLVMs) by coupling **<u>b</u>**iomass **<u>e</u>**stimation and model inference in an **<u>e</u>**xpectation-**<u>m</u>**aximization-like algorithm (BEEM). Surprisingly, BEEM outperforms state-of-the-art methods for inferring gLVMs, while simultaneously eliminating the need for additional experimental biomass data as input. BEEM's application to previously inaccessible public datasets (due to the lack of biomass data) allowed us for the first time to construct ecological models of microbial communities in the human gut on a per individual basis, revealing personalised dynamics and keystone species.

## 1. Introduction

A growing body of literature points to the important roles that different microbial communities play in diverse natural environments^1,2^ and the human body^3^. This has particularly been aided by advances in next-generation sequencing technology, allowing for rapid, cost-effective taxonomic and functional profiling, combined with computational analysis that has helped associate the state of the microbiome with various environmental conditions^1,4^ and human diseases^5–8^. Microbiomes are also constantly evolving and there is now a growing appreciation that complex interactions between community members^9,10^ shape community dynamics^11,12^ as well as overall function^13,14^. A systems view of the microbiome is thus essential for understanding and rationally manipulating it^15^.

Because of its importance, there have been many approaches proposed to study microbial interactions and dynamics. Experimental approaches have ranged from simple two species co-culture experiments^16–18^, all the way to complex, multi-stage reactor models^19^. Analytical approaches^20^ frequently use simple correlations between the abundances of various taxa in cross-sectional datasets to infer microbial interactions^21–23^. There are several challenges that need to be addressed in such analysis including the compositionality of sequencing data^21–24^, low sensitivity and specificity of such methods^25,26^, and the inability to infer directionality of interactions or dynamics of the system^20^.

The most commonly used approach for modeling microbial ecology is based on classical predator-prey systems, also referred to as generalized Lotka-Volterra models (gLVMs). gLVMs are based on ordinary differential equations (ODE) that model the logistic growth of species, naturally capture predator-prey, amensalistic and competitive interactions, and have been applied to study dynamics of microbial ecosystems ranging from simple communities on cheese^27,28^ to the human microbiome^15,26,29–32^. More importantly, from a practical perspective, gLVMs have been used for a range of applications including identifying potential probiotics against pathogens^15,29,30^, forecasting changes in microbial density, characterizing important community members (e.g. keystone species^26^) and to analyze community stability^30,32,33^.

Despite this, a key limitation of gLVMs that restricts applicability and wider use is the requirement for microbial abundance data on an absolute scale. Microbiome analysis using high-throughput sequencing naturally provides relative abundance estimates with what is often referred to as “compositionality bias”^21,22,24^, and cannot be directly used to estimate gLVM parameters^31^. Scaling relative abundances to an absolute scale typically requires additional experimental data that is either not readily available (as is true for the vast proportion of publicly available datasets), is technically challenging to directly quantitate for different sample matrices and complex communities (e.g. using flow cytometry^34,35^), or can suffer from significant technical^36–38^ and biological noise^39^ (e.g. using 16S rRNA qPCR^15^,^29^,^30^).

In the face of these technical challenges, gLVM inference can seem daunting, especially because relative abundances do not seem to carry any information related to absolute scale. Remarkably, we show that suitable scaling factors can be directly inferred from metagenome sequencing data, through an algorithm that also imposes constraints based on gLVM inference (BEEM). This is achieved based on an expectation-maximization-like approach^40^ that alternates between learning scaling factors and gLVM parameters, and thus obviates the need for experimental scaling factors which otherwise limit the use of many existing datasets. Based on synthetic data where biomass is precisely known, we show that BEEM estimated gLVM parameters are as accurate as those estimated with true biomass values, and significantly more accurate than what could be expected with commonly used (16S rRNA based) experimentally determined biomass estimates. Using data from a freshwater microbial community with flow cytometry based gold-standard cell counts, we show that biomass estimated using BEEM has good concordance with the gold-standard and improves significantly over existing techniques to normalize data. Leveraging BEEM’s unique ability to learn gLVMs from relative abundance data, we analyzed publicly available datasets that represent the longest human gut microbiome time-series data available to-date^41–43^. This analysis highlighted the personalized dynamics of gut microbial biomass in different individuals, with communities driven by distinct interaction networks and hub species. Our analysis suggests an emergent model for gut microbial dynamics where relatively low abundance species may play key roles in maintaining gut homeostasis.

## 2. Results

### 2.1 Experimentally obtained biomass estimates can lead to inaccurate gLVMs

The gLV equations model the growth rate 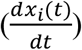 of each microbial species *i* as a function of absolute densities (*x_i_*(*t*)) of all the *p* species in a community:
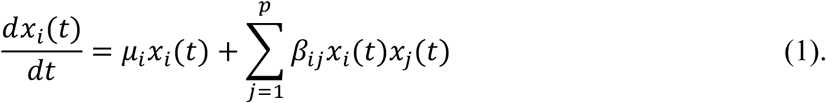

In the above model, the intrinsic growth rate parameter (*μ_i_*) and self-interaction parameters (*β_ii_*) define the logistic growth behavior of species *i*. In addition, the model also captures the impact of the absolute density of species *j* on the growth rate of species *i* through additional parameters (*β_ij_*, *i* ≠ *j*), assuming a linear and additive effects model. As high-throughput sequencing based approaches to analyze microbiomes only provide relative abundance estimates, scaling factors related to the total biomass for each sample are then needed to accurately fit gLVMs in practice.

The predominantly used approach to estimate total biomass is to quantify copy number of the 16S rRNA gene using quantitative PCR (qPCR)^15,29,30^. However, 16S qPCR estimates have been reported to have high technical noise, with a coefficient of variation (CV) ranging from 11% to 75%^36–38^. To reconfirm this, we reanalyzed 16S qPCR data from a recent microbiome modeling study on *C. difficile* infections^30^ and observed low concordance across technical replicates (Spearman *ρ*<0.22; **Figure 1A** and **Supplementary Figure 1A**), as well as high coefficient of variation (mean CV=51%). Another critical source of error with 16S qPCR based biomass estimates is biological, and arises due to the fact that bacteria can have widely varying number of copies of the 16S rRNA gene, even within the same ecological niche. For example, the 16S gene copy number of the four major gut bacterial phyla cover a broad spectrum (**Figure 1B**), ranging from a single copy to 15 copies^39^. Correspondingly, 16S qPCR estimated biomass of a community dominated by *Firmicutes* can be twice as much as that of a community dominated by *Bacteriodetes*, even if both communities have exactly the same cell density. Such large relative errors (~100%) can then have a significant impact on the accuracy of gLVMs estimated from the data, as we show below.

**Figure 1.**
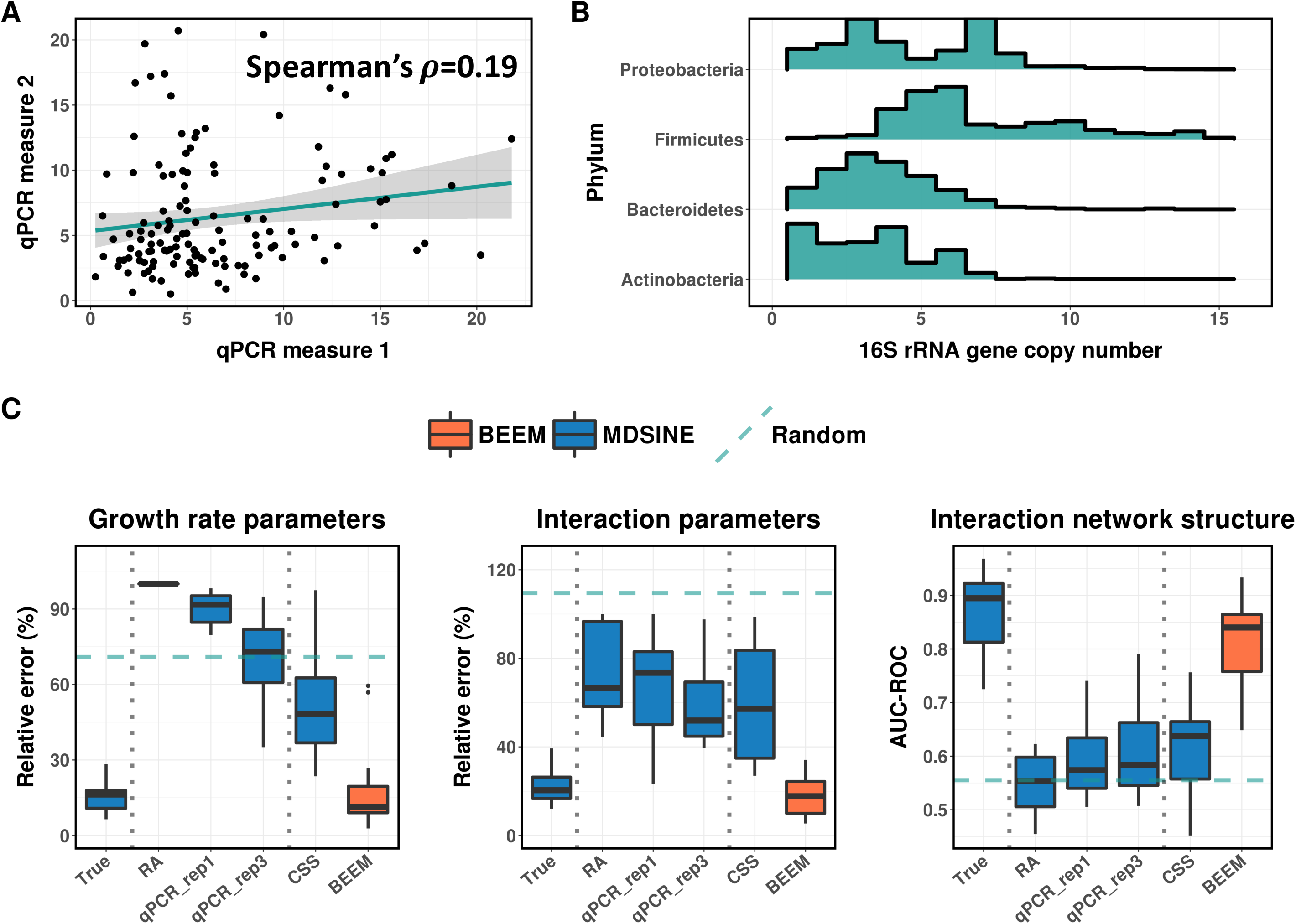
Noise in experimentally determined biomass severely distorts gLVM parameter estimation. (A) Scatter plot with fitted linear regression line for two 16S qPCR technical replicates from Bucci *et al*. (B) Copy number variation for 16S rRNA genes in members of four major phyla of human gut bacteria. (C) Relative impact of different experimental (qPCR_rep1 – 1 qPCR replicate, qPCR_rep3 – mean of 3 qPCR replicates) and computational (RA – relative abundance, CSS – CSS normalization) data scaling approaches on gLVM parameter estimation (BVS algorithm for MDSINE), in comparison to using true biomass or using BEEM. Boxplots represent the summary of 15 simulations (10 species, 30 replicates with 30 time points each) and three different metrics are shown here including median relative error for growth rate (***μ***) and interaction (***β***) parameters, and AUC-ROC for the interaction network. Dashed horizontal lines represent the performance of randomly generated parameters from the simulation model.

To test the impact of biomass estimation errors on model inference, we generated synthetic datasets (10 species community) based on parameters inferred from real datasets, similar to the approach in Bucci *et al*^29^ (see **Materials and Methods**). This framework allows us to carefully evaluate the impact of different levels of noise in a setting where model parameters are known. We noted that, given error-free biomass data, a state-of-the-art method (MDSINE^29^) was able to infer model parameters with median relative error <20% and with ~90% median AUC-ROC (area under the sensitive-specificity tradeoff curve) for interaction terms (*β*; **Figure 1C**, True). However, as expected^31^, directly using relative abundance estimates without scaling them increased median relative error for parameter estimates to >60% (**Figure 1C**, RA), with AUC-ROC for interaction terms being comparable to randomly generated parameters from the prior model for the simulation (**Figure 1C**, Random). Similar performance was obtained using another model fitting algorithm that works with relative abundance data and assumes small fluctuations in biomass values (LIMITS^26,44^; **Supplementary Figure 1B**). Using biomass estimates with error profile similar to real qPCR data (CV=51%; without systematic errors due to varying copy number of the 16S rRNA gene), surprisingly, did not improve performance substantially when one technical replicate was provided (**Figure 1C**, qPCR_rep1), and even with three technical replicates, growth rate parameter estimates (median relative error >70%) were comparable to random (**Figure 1C**, qPCR_rep3). These results highlight that experimental errors in biomass estimates can significantly impact gLVM parameter estimation even in a relatively well-controlled setting where model assumptions are strictly applied.

### 2.2 Joint estimation of biomass and model parameters with BEEM

In order to address the challenges of noisy experimental biomass data and, in general, to make gLVM modeling more widely applicable where biomass estimates are not available, we explored the idea of learning gLVM parameters directly from relative abundance data. To achieve this, we first note that model equation 1 can be expressed in terms of relative growth rates by dividing both sides of the equation by *x_i_*(*t*):
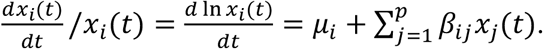

By explicitly introducing relative abundances 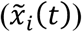 and total biomass (*m*(*t*), where 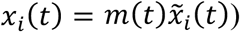, we get:
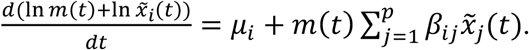

The biomass terms on the left-hand-side of the equation can be eliminated by subtracting the equation of a selected species *r* from the equations for all other species, resulting in a new system:
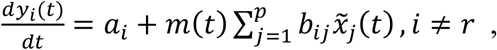

where 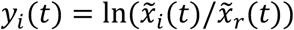 and the equations are re-parameterized by *a_i_* and *b_ij_*, which are related to the original parameters (*a_t_ = μ_i_ − μ_r_* and *b_ij_ = β_ij_ − β_rj_*). This new system has the advantage that all unknowns are on the right-hand-side of the equation and the gradient term on the left-hand-side can be estimated directly from relative abundance data through spline smoothing and numerical differentiation^15,26,29,30^.

We then made the observation that the above equations can be re-written as two regression problems across two dimensions of the data matrix 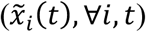:

1. For each species *i*, the corresponding parameters *a_i_* and *b_ij_* can be solved through gradient matching^15,26,29,30^, given the biomass at each time point *t* (*m*(*t*)).
2. For each time point *t*, the biomass can be solved for via regression given the model parameters ***a*** and ***b*** for all the species.

The interlock of the above two problems provides the basis for an expectation-maximization-like algorithm that alternates between estimating model parameters and biomass iteratively and forms the core of BEEM (see **Materials and Methods** for details). Note that estimates provided by BEEM for the biomass act as scaling factors to bring abundances across species and time points to the same scale for learning gLVMs.

On the synthetic datasets used in section 1, we noted that despite not having any biomass data to work with, BEEM was a significant improvement over naïve analysis based on relative abundance data, as well as results based on scaled relative abundances with noisy biomass data (~3× reduction in relative error; **Figure 1C**, BEEM). In fact, BEEM estimated parameters were nearly as accurate as those obtained using noise-free biomass data (relative error for growth rate and interaction terms), except for a slight decrease in AUC-ROC for interaction terms (primarily due to rounding errors that provide non-zero estimates for zero terms). In comparison, other competing approaches (RA, qPCR, CSS) provided AUC-ROC performance similar to what is expected at random. Normalization approaches such as CSS^45^ and TMM^46^ (**Figure 1C**, CSS; **Supplementary Figure 1B**; **Materials and Methods**) were tested here as control analytical methods, but are not expected to work in general as they are designed to identify scaling factors that do not change across samples. We noted that BEEM’s significant improvement over other experimental and computational approaches, and its ability to closely approximate analysis using true biomass estimates is a robust feature that remains valid even when experimental biomass estimates are significantly better (CV=5%, as expected from flow-cytometry data) and while using different parameter estimation approaches (**Supplementary Figure 1B**).

### 2.3 BEEM accurately estimates gLVM parameters and biomass in diverse model settings

As in any situation where parameters have to be estimated, a sufficient number of data points (multiple biological replicates) are needed to get accurate gLVM models and this in turn impacts BEEM’s biomass estimates. In order to further study BEEM’s performance characteristics, we generated synthetic datasets with varying number of species and data points, comparing BEEM’s results to those obtained with noise-free biomass data and the same gradient matching algorithm (BLASSO, see **Materials and Methods**) as used internally in BEEM. As expected, when the number of species increases but the number of data points remains constant (60 replicates with 30 timepoints), gLVM parameter estimation becomes harder (**Figure 2A**). However, despite the quadratic increase in the number of parameters, performance for both BLASSO (with true biomass) and BEEM seems to only degrade linearly (**Figure 2A**). In addition, even when the model has 25 species (650 model parameters) and can thus capture over 90% of the human gut microbiome^47^ (Supplementary Figure 2), interaction parameters estimated by BEEM were nearly as accurate as those with true biomass (**Figure 2A**), though growth rate parameters were more affected. We also noted that median relative error for biomass estimates from BEEM was generally well controlled (<10%; **Figure 2B**).

**Figure 2.**
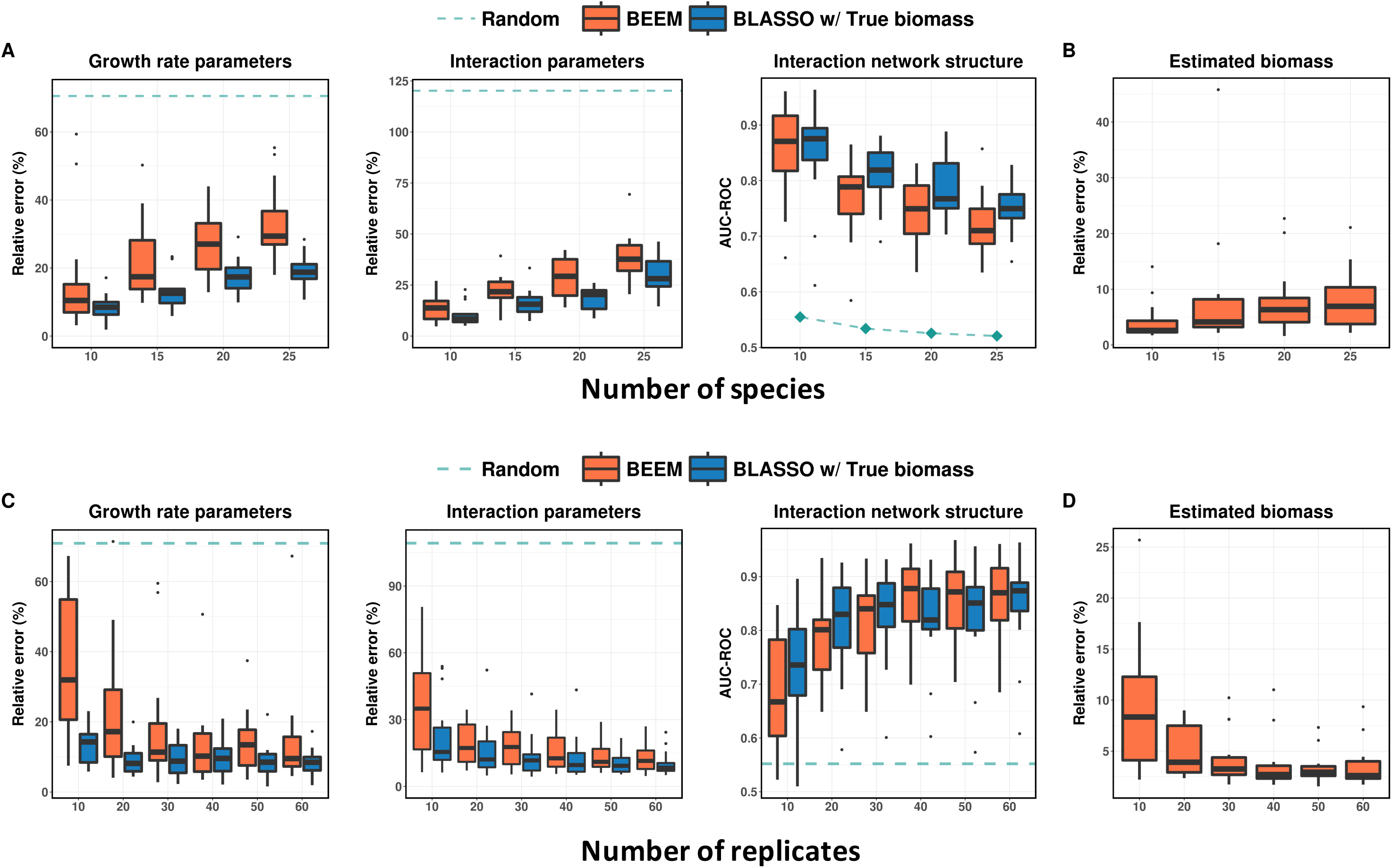
Robustness of parameter estimation with BEEM. (A) Results with increasing number of species but fixed number of replicates (50). As expected, parameter estimation gets harder but BEEM’s performance tracks the ideal case using BLASSO with true biomass values, especially for interaction parameters. (B) Median relative error in biomass estimates remains less than 10%. (C) Results with increasing number of replicates and fixed number of species (15). BEEM’s performance converges to that of BLASSO with true biomass as the number of replicates increases. (D) Median relative error in biomass estimates reduces noticeably as the number of replicates increases.

Increasing the number of data points available for model fitting for a fixed number of species (10) improved performance for both BLASSO with true biomass and BEEM, as expected. Performance improvements were most notable when going from 10 to 20 replicates and plateaued out after that (30 timepoints; **Figure 2C**). In general, after 20 replicates, differences between BLASSO and BEEM were small, especially in terms of estimating interaction parameters. Similarly, biomass estimates from BEEM had median relative error <5% when 20 replicates were available (**Figure 2D**). In general, our analysis suggests that inherent limitations in gradient matching based on estimated gradients from data were a greater source of error for gLVM parameter estimation in many of our experiments than errors in BEEM estimated biomass values.

To assess BEEM’s performance for biomass inference in real-world datasets we analyzed data from a recently published study on freshwater microbial communities^34,35^, which to our knowledge is the only one to have longitudinal metagenome sequencing data as well as flow-cytometry based gold-standard biomass estimation. Notably, the flow cytometry data in this study was reported to have high reproducibility (CV<5%)^34^, and therefore was suitable for use as the ground truth for total biomass. Surprisingly, with only 57 time points in total across two replicate experiments, BEEM was able to infer the total biomass for a 26-species community accurately solely based on 16S sequencing based relative abundances. BEEM estimated biomass values showed strong correlation with flow cytometry data (BEEM: Spearman’s ρ=0.72; **Figure 3A**) and its trajectories closely tracked measured fluctuations (**Figure 3B**). In contrast and as expected, normalization approaches provided estimates that had either weak correlation (CSS: Spearman’s ρ=0.36) or negative correlation with experimentally determined values (TMM: Spearman’s ρ=-0.11; **Figure 3A**).

**Figure 3.**
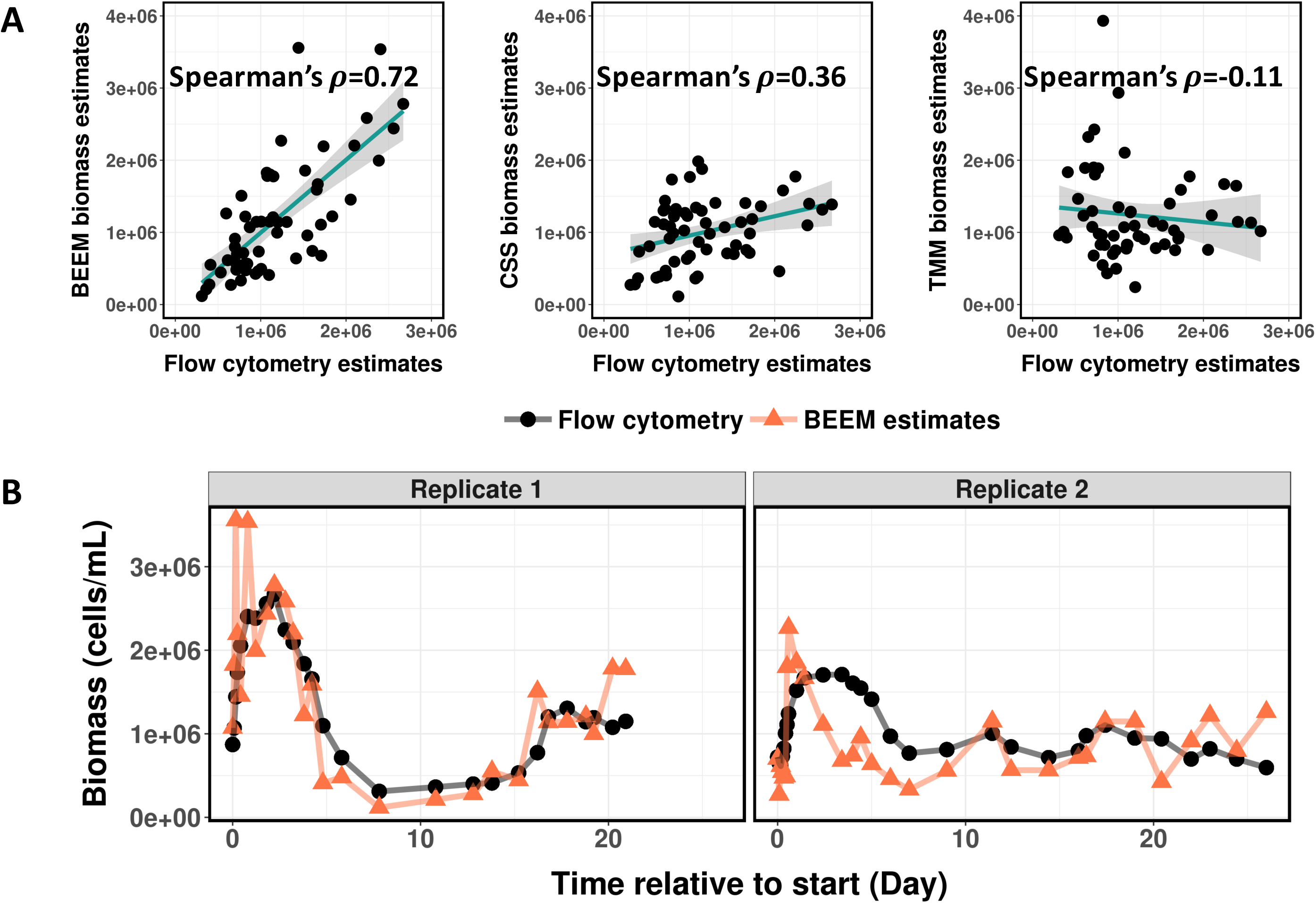
Concordance of BEEM estimated biomass with gold standard experimental measurements. (A) Scatter plots with fitted linear regression line highlighting that BEEM’s biomass estimates are notably more concordant with flow cytometry based values compared to CSS and TMM normalization based estimates. (B) BEEM estimated biomass values (orange) compared to gold standard measurements using flow cytometry (black).

### 2.4 Personalized gut microbial dynamics and keystone species

The development of BEEM allows us to analyze previously generated datasets in a gLVM framework, even when biomass measurements were not made in the original study. To showcase this capability, we applied BEEM to the longest (over one year) and most densely (almost daily) sampled human gut microbiome time-series datasets available to date (four individuals: DA, DB from David *et al*^42^ and M3, F4 from Caporaso *et al*^41^). BEEM estimated models exhibited a good fit to the data, with predicted relative abundances for a day based on numerical integration from the previous day being in high concordance with observed data (median Spearman’s ρ = 0.83). As BEEM directly infers daily biomass values, we plotted these and observed distinct individual-specific patterns: while subject DA’s biomass was found to vary relatively smoothly, following an approximately cyclic pattern with a period of about three months (**Figure 4A**), subject M3’s biomass fluctuated to a greater extent on a day to day basis with no clear trend (**Figure 4B**). Similar patterns were observed in parts for subjects DB and F4, which had a greater resemblance to DA overall (**Supplementary Figure 3A, B**). The fluctuations predicted in M3’s biomass were also found to be accompanied by frequent blooms of rare taxa that were otherwise not detected at other time points^43^ and may be a consequence of this instability in the community. In contrast, the smoother progression of DA’s biomass may be a reflection of the relative stability of the gut community in this individual, though the source of the observed cyclic patterns deserves to be explored further. As an initial hint, we noted that the strongest association between DA’s biomass and reported metadata was a negative correlation with calcium intake (Supplementary Figure 4).

**Figure 4.**
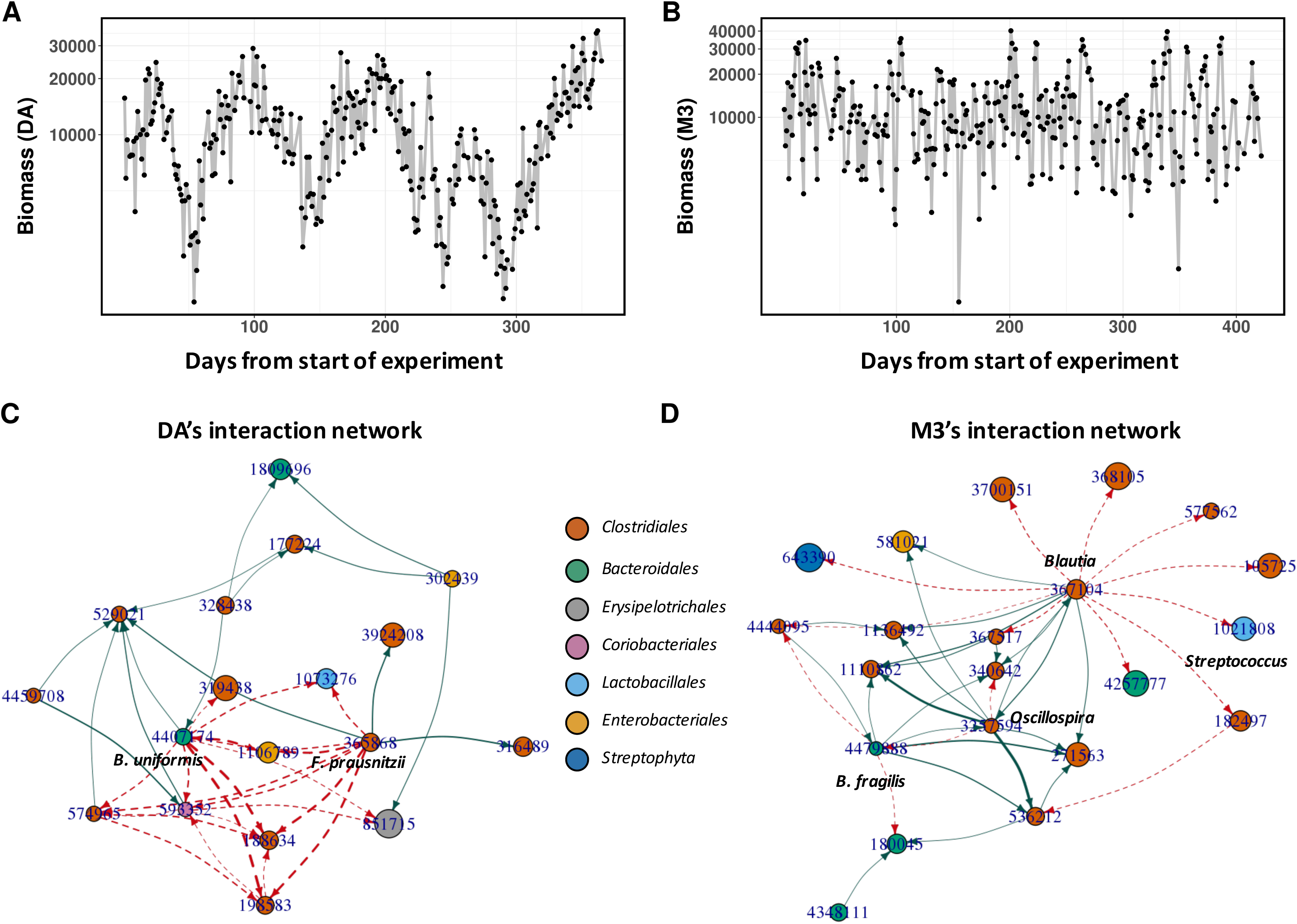
BEEM analysis of year long gut microbial time-series datasets. (A, B) BEEM estimated biomass values for two individuals (DA and M3) with daily sampled, year long gut microbial time-series datasets from David *et al*^42^ and Caporaso *et al*^41^. Interestingly, while M3’s biomass fluctuates rapidly, DA’s biomass seems to vary in a more defined fashion with a periodicity of around 3 months. (C, D) Graphs representing non-zero interaction terms in gLVM models learnt individually for DA and M3 using BEEM. Dashed and solid edges represent positive and negative interactions respectively. Edge widths are proportional to the interaction strength and node sizes are proportional to the log-transformed mean relative abundance of the corresponding species. Nodes are labeled with GreenGenes IDs and colored according to order level taxonomic annotations.

Concordant with their distinct biomass dynamics, DA and M3 also exhibited microbial interaction networks that were unique to them (**Figure 4C, D**). DA’s network was defined by hub nodes for *Feacalibacterium prausnitzii* and *Bacteroides uniformis*, two species with many beneficial roles and frequent associations with a healthy gut^48,49^. The hubs were found to negatively affect the growth of *Enterobacteriaceae* species, consistent with previous reports for *B. uniformis*^50^ and *F. prausnitzii*^51–53^. In comparison, the major hub nodes in M3’s network were a *Blautia* and an *Oscillospira* species that were connected by a positive feed-forward loop. Additionally, we found that abundances of the *Blautia* and *Oscillospira* species were significantly negatively correlated with total biomass in M3’s gut microbiome (**Supplementary Figure 5**). Feed-forward loops have been implicated in destabilizing effects on ecosystems^32^ and so these observations may explain the unstable behavior of M3’s biomass as well as the corresponding susceptibility to invasive blooms of rare taxa^43^. *Oscillospira’’* s protective role in M3’s gut flora is further indicated by its parasitic relationship (negative-positive loop) with another hub species *B. fragilis*, an opportunistic pathogen that has been associated with diarrhea^54^. Interestingly, several of the transient species in M3’s gut microbiome were observed to be at the periphery of the network, with a single incoming edge indicating that their abundances were being influenced by a hub species. For example, this was observed for several *Streptococcus* species that are primarily oral commensals and could be transient colonizers of the gut^55,56^.

Despite differences in the identity of species in their interaction networks, the various individual-specific networks shared some common features, including the presence of a few hub nodes that negatively influenced many other species, and were generally not the most abundant species in the community (**Figure 4C, D** and **Supplementary Figure 3C, D**). Overall, we also found that the ratio between out- and in-degree of species in the networks were negatively correlated with their median relative abundances (**Supplementary Figure 6**), suggesting that the hub species in the interaction network, that are often considered as keystone species for the community^26,57^, are typically not the abundant species in the gut microbiome. We further confirmed this observation by analyzing a large collection (840 healthy individuals) of gut microbiome datasets^47^, to find that the core species in the gut microbiome were also frequently not the most abundant species (Supplementary Figure 7). Together, these observations suggest a model for the gut microbiome where relatively less abundant species in the community are more stable colonizers of the host, and by virtue of their impact on the growth of other species in the community, play an important role in defining its dynamics in different individuals.

## 3. Discussion

A major limitation of most microbiome profiling datasets available to date is the restriction to relative abundances and the ‘compositionality’ of this data has led to significant challenges even when performing common statistical tests for correlated abundances^58^. These issues are amplified when considering systems models such as gLVMs, and our analysis here confirms that model parameter estimates can be severely distorted if relative abundances are not correctly scaled. In ecological models such as gLVMs, interactions between species are naturally a function of the absolute density of species in a community rather than their relative abundances. Correspondingly, while autoregression based methods such as sVar^43^ and ARIMA^59^ provide an alternative for model fitting with relative abundance data, their models and parameters are not ecologically interpretable. In addition, experimental approaches to measure scaling factors are generally seen as a laborious and occasionally feasible way to work with absolute abundances, but as we show here, this may not be the case if care is not taken to ensure that experimental noise is minimized and sufficient number of replicates are analyzed. By eliminating the need for additional experimental data, BEEM greatly expands the applicability of gLVMs to microbiome datasets, and its robustness could simultaneously improve the quality of models and scaling factor estimates, as observed in our synthetic and real datasets. Explicitly modelling microbial interactions through gLVMs has proven to be a powerful framework for studying microbial community dynamics^15,26–32^, and the approach used in BEEM could also be extended (with minimal modifications) to time-series with external perturbations (e.g. antibiotics usage)^15,29,30^, as well as systems models for gene expression regulation based on RNA-seq data^60^.

Due to limited availability of absolute abundance data, gLVMs have generally been constructed by aggregating information across experiments and individuals^15,29,30^. We exploited the availability of year-long time series datasets and BEEM’s facility with relative abundances to construct individual specific gut microbiome gLVMs for the first time. Intriguingly, we observed that our inferred scaling factors suggest that gut microbial biomass has distinct dynamics across different individuals. Consistent with a recent study on 20 individuals where human gut microbial biomass (measured via flow-cytometry) was found to have high variation (CV ≈ 53% within a week)^58^, we also noted high variability over time across the four individuals we analyzed (CV ranging from 49% to 76% over a year). Additionally, we observed cyclic behavior of biomass trajectories in multiple individuals, similar to the seasonal patterns reported in hunter-gatherers of western Tanzania^61^, and the conserved patterns observed in other mammals across evolutionary timescales^62^. Similar patterns have not been reported before for western city dwellers, perhaps due to the confounding effects of aggregate analysis across individuals and the impact of highly diverse diets. BEEM analysis, however, suggests that the underlying patterns may still be conserved in urban subjects and may be more general than previously believed.

Our inference of gLVM models for each individual allows us to identify specific microbial species and the kinds of interactions that they have, to account for the distinct dynamics that were observed. For example, the positive feed forward loop observed between the hubs in M3’s gut microbiome provides a specific, plausible and testable hypothesis to explain the instability observed there, and this capability can be valuable in future studies where targeted interventions are feasible. Despite differences in the microbial interaction networks observed for different individuals, a shared feature seems to be the presence of relatively lowly abundant species that act as hub nodes in the network. A similar pattern was seen in cross-sectional data as well where frequently shared “core” gut microbiome species tend to not be the most abundant species in the community. These observations point to a model where species at low relative abundances stably colonize the gut (e.g. mucosa-associated ones) compared to abundant but transient (lumen-associated) bacteria, and play an important role in defining gut microbiome dynamics. In particular, hub species were frequently found to negatively regulate more transient species in the community, in agreement with the known role of mucosa-associated species in providing colonization resistance against invasive pathogenic species^63^.

An important point that we noted in the gut microbiome datasets that were analyzed here is the limited number of stable species (prevalent in most time points for an individual) that are shared across individuals. This feature makes it infeasible to learn gLVM models by merging short time-series datasets across different individuals. Similar constraints might be present in other microbial communities as well, including specific challenges in measuring total biomass in complex matrices^58^, and thus the development of BEEM makes it more feasible to generate the long and densely sampled datasets that are needed for such models. In addition, the analysis in BEEM can potentially be directly extended to cross-sectional datasets if the corresponding communities are believed to be at equilibrium (i.e. 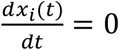, for all species). This extension would significantly expand the amount of data that could be used and thus allow us to learn even more complex models in the future. As is the case for any modelling approach, no model is expected to be perfect, but as they capture more and more features of real systems, we can expect that their predictions become increasingly useful. BEEM’s development therefore serves as an important step in expanding the use of modelling approaches to study microbial community dynamics and rationally identify appropriate perturbations.

## 4. Materials and Methods

### 4.1 BEEM’s core algorithm

As introduced in **Section 2.2**, the gLVM model in equation 1 can be first simplified by dividing *x_i_*(*t*) on each side, and then re-written in terms of total biomass *m*(*t*) (i.e. 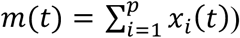) and relative abundances 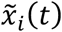 (i.e. 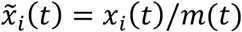) as shown below:
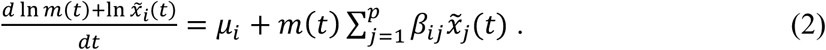

To eliminate the biomass related term in the left-hand-side of the equation, we subtract the corresponding equation for a reference species *r* (species with lowest CV, by default) from both sides of the system, resulting in additive log ratio (ALR) transformed^64^ relative abundances 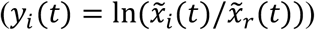 on the left-hand-side and a re-parameterized right-hand-side:
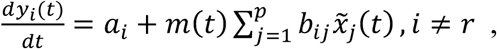

where *a_i_ = μ_i_ − μ_r_* and *b_ij_ = β_ij_ − β_rj_*.

An estimate for *dy_i_*(*t*)*/dt*, denoted as *Y_it_*, can be calculated as the derivative of a piece-wise polynomial spline fitted to the ALR transformed relative abundances (*y_i_*(*t*), see **Section 4.2** for details). BEEM then estimates the model parameters ***a, b*** and the biomass ***m*** using an EM-like algorithm with the following sum of squared error objective function:
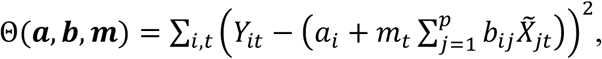

where 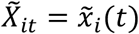 and *m_t_ = m*(*t*) are the variables written in their matrix representations.

The EM-like algorithm in BEEM works by iterating two steps, an E-step and an M-step, to convergence as detailed below:

**Model parameter estimation with Bayesian lasso (E-step):** In iteration *T*, with estimated biomass from the previous iteration 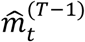, BEEM estimates 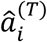 and 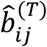 for each *i* (*i ≠ r*) based on the following regression problem (also known as gradient matching):
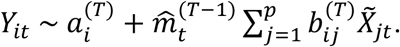

Solving the above system is often limited by the amount of data available in practice. For microbial communities, it is usually assumed that the interaction vector (*β_ij_*) is sparse (i.e. a species is only directly affected by a small number of other species). Consequently, the transformed matrix *b_ij_* is also sparse and BEEM estimates it using a sparse regression technique based on a Bayesian approach (Bayesian lasso - BLASSO^30^; R package “monomvn” version 1.9-7; default parameters)^65^.

**Biomass estimation with linear regression (M-step):** With 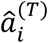 and 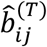 from the E-step, the biomass 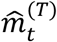 for each *T* can be computed as the coefficient of the following linear regression:
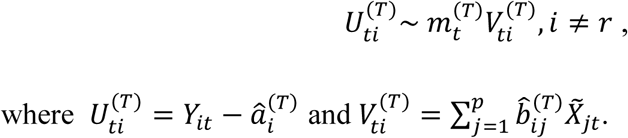

**Initialization:** For the initialization step in its EM-like algorithm, BEEM assumes that scaling factors inferred from a commonly used normalization approach for metagenomic data (Cumulative Sum Scaling - CSS^66^) provides a reasonable starting point for the algorithm to then learn better scaling factors. Note that, as expected, scaling factors from CSS normalization and BEEM cannot recapitulate the absolute scale corresponding to experimental measurements (e.g. by qPCR or flow cytometry), and so their estimates were scaled to the same median value across the time series as experimental measurements for subsequent comparisons.

**Termination and parameter estimation:** The E- and M-step in BEEM are run until convergence or a user specified maximal number of iterations. The search was assumed to have reached convergence (to a local optimum) when the mean squared error (MSE) for the M-step starts to increase by more than 10% compared to the minimal MSE observed. In practice, on the datasets analyzed in this study, convergence takes a few hours using 4 CPUs. Estimates for 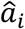, 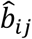 and 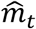 were calculated as the median of the values from all iterations whose MSE was within 10% of the minimal MSE.

### 4.2 Robust parameter estimation with BEEM

In our experiments with synthetic and real data, we noted that gLVM modelling can be sensitive to noise and outliers in the data, and this in turn could affect estimation of scaling factors with BEEM. To address this, we refined the core algorithm in BEEM with additional pre-processing steps that further enable robust parameter estimation.

**Outliers in relative abundance data:** We observed in our numerical analysis that outliers in the abundance data could notably affect the spline fitting procedure and lead to spurious gradient estimates. To obtain more robust spline fitting, an over-smoothed spline was first fitted to *y_i_*(*t*) (function “smooth.Pspline” from R package “pspline”^67^ with maximal degree of five and a large smoothing parameter “spar=1e10”) to calculate the absolute error in fitted values (*e_it_* = |*y_i_*(*t*) − *y_i_*(*t*)^smoothed^|), and points with absolute error larger than expected (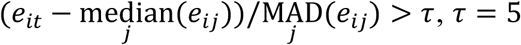 by default) were then filtered out. The final smoothing spline was fitted (degree of five and smoothing parameter selected using cross validation) to the remaining data to calculate the estimated gradients *Y_it_*. In addition, outliers in biomass estimated from the previous iteration 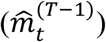 were identified in the same way and replaced with interpolated values from the spline.

**Outliers in estimated gradients:** In practice, gradient matching based methods (including the various algorithms implemented in MDSINE) were found to be sensitive to outliers in the estimated gradients (i.e. *Y_it_*). To identify outliers in a time series (*Y_it_*, for all *t*) a local regression (LOESS) smoother was fitted to de-trend *Y_it_*, and the outliers were filtered out as described above.

**Estimating constrained biomass values:** For each time point, biomass was estimated as the slope of a linear regression (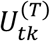 against 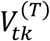) where outliers in both 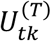 and 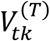 were identified and removed following a standard boxplot approach i.e. as deviations from the median by more than 1.5 × inter-quartile range. In addition, the biomass was constrained to be positive by removing points where 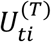 and 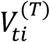 had different signs.

### 4.3 Recovering gLVM parameters

Based on the previously stated assumption that the interaction matrix ***β*** is sparse, most entries in each column are expected to be zero and thus the median value for the *j*^th^ column in ***b*** would be expected to be −*β_rj_*, allowing us to infer back all the other rows of ***β*** (*β_ij_* = *b_ij_ + β_rj_*). BEEM then assigns a Z-score like confidence value (*s_ij_*) to each entry of ***β***, by dividing the estimated interaction strength by the column standard deviation 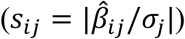. The growth rate vector ***μ*** is not expected to be sparse but can be recovered by directly solving the original gLVM system (equation 2), using the already derived estimates for scaling factors and ***β***. For robustness, BEEM estimates the growth rate for each species as the median of positive estimates across all time points.

### 4.4 Datasets and evaluation metrics

**Simulated datasets:** MDSINE’s Bayesian variable selection (BVS) algorithm (with spline smoothing option and minor bug fixes: https://bitbucket.org/chenhao_li/mdsine) was used to estimate parameters from the *C. difficile* infection dataset provided with the package^30^. Simulated datasets were then generated based on these estimated parameters following the procedure described in Bucci *et al*^30^ (excluding perturbations). Noisy abundances were obtained by sampling from Poisson distributions^68^ with means based on scaled abundances at each time point (sum = 5×10^4^). Simulated qPCR and flow cytometry based values for total biomass were generated from log normal distributions with coefficients of variation (CV) that matched those seen in real datasets (qPCR=51%^30^, flow cytometry=5%^34,35^).

**Dataset from Props et al:** The original OTU table was obtained from the authors^35^. Samples for the “operation” stage, where the environment had roughly constant temperature were selected for BEEM analysis. OTUs with low mean relative abundances (<0.1%) were excluded, resulting in 26 OTUs across 58 time points from two replicates.

**Dataset from Gibbons et al:** This dataset included four long time series collected by David *et al*^42^ and Caporaso *et al*^41^. The original OTU tables^43^ were filtered to keep only top OTUs based on prevalence (>10 reads in most of the samples). In total, 26 and 22 OTUs were left for samples from David *et al* and Caporaso *et al*, respectively. In order to assess the robustness of the inferred network, BEEM was run with 30 different seeds and edges with confidence score *s_ij_* ≤ 1 in more than 50% of the networks were kept. The final biomass was obtained by taking the geometric mean across all 30 runs (**Supplementary File 1**).

**Metrics for evaluation:** The following metrics were used for evaluating inference algorithms:

1. Median relative error (MRE) for estimates 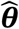 when the true values are ***θ***: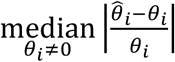.
2. Area under receiver operating characteristic curve (AUC-ROC) for the inferred microbial interactions: Absolute values of parameters was used to rank predicted edges for BLASSO and LIMITS (implemented in R package seqtime_0.1.1^44^, default parameters), while confidence scores were used for Bayesian Variable Selection (BVS) in MDSINE and for BEEM.

### 4.5 Software and reproducibility of results

BEEM is available under the MIT license and can be downloaded from https://github.com/csb5/BEEM. Scripts to reproduce the results presented in this work are also available at this website.

## Acknowledgements

We would like to thank Dr. Lawrence David, Dr. Eric Alm, Dr. Ruben Props and Dr. Nico Boon for sharing OTU tables from their studies^35,42^.

## Funding

This work was supported by funding to the Genome Institute of Singapore from the Agency for Science, Technology and Research (A*STAR), Singapore.

